# Predicting functional neuroanatomical maps from fusing brain networks with genetic information

**DOI:** 10.1101/070037

**Authors:** Florian Ganglberger, Joanna Kaczanowska, Josef M. Penninger, Andreas Hess, Katja Bühler, Wulf Haubensak

**Affiliations:** VRVis Research Center, Donau-City Strasse 1, 1220 Vienna, Austria; Research Institute of Molecular Pathology (IMP), Vienna Biocenter (VBC), Dr. Bohr-Gasse 7, 1030 Vienna, Austria; Institute of Molecular Biotechnology of the Austrian Academy of Sciences (IMBA), Vienna Biocenter (VBC), 1030 Vienna, Austria; Institute of Experimental and Clinical Pharmacology and Toxicology, Friedrich-Alexander University Erlangen-Nuremberg, Fahrstrasse 17, 91054 Erlangen, Germany

## Abstract

A central aim, from basic neuroscience to psychiatry, is to resolve how genes control brain circuitry and behavior. This is experimentally hard, since most brain functions and behaviors are controlled by multiple genes. In low throughput, one gene at a time, experiments, it is therefore difficult to delineate the neural circuitry through which these sets of genes express their behavioral effects. The increasing amount of publicly available brain and genetic data offers a rich source that could be mined to address this problem computationally. However, most computational approaches are not tailored to reflect functional synergies in brain circuitry accumulating within sets of genes. Here, we developed an algorithm that fuses gene expression and connectivity data with functional genetic meta data and exploits such cumulative effects to predict neuroanatomical maps for multigenic functions. These maps recapture known functional anatomical annotations from literature and functional MRI data. When applied to meta data from mouse QTLs and human neuropsychiatric databases, our method predicts functional maps underlying behavioral or psychiatric traits. We show that it is possible to predict functional neuroanatomy from mouse and human genetic meta data and provide a discovery tool for high throughput functional exploration of brain anatomy *in silico*.

## Introduction

The wealth of data from brain initiatives and the increasing amount of functional genetic information creates opportunities to mine these resources for insights into the genetic and neuronal organization of brain function and behavior. Recent studies correlated brain gene expression maps with structural information to enhance our understanding of genetic and anatomical parcellations of the brain (1, 2) and its functional networks (3). These studies have been used, for instance, to explore development and physiological regulation of structural connectivity and extract functional networks *in silico* (Supplementary Note 1). Collectively, these results suggest that functional genetic information, brain gene expression data and connectomes can be successfully used for functional exploration of the brain (Supplementary Fig. 1).

Here, we mine these resources to understand how genes control behavior. A major challenge in this regard is that behaviors are inherently multigenic and, consequently, identifying the neural networks through which these gene sets interact to express that function is not trivial. Discovery tools that give computational predictions would provide an ideal entry point into this problem.

Most established approaches that map genetic information to brain data relate gene co-expression correlation of functionally grouped genes with structural connectivity (2–5). Correlative analysis primarily dissects brain organization based on the similarities of regional gene expression (Supplementary Note 1). It primarily reflects transcriptomic similarities, globally or for subsets of genes, but it is not tailored to directly predict functional synergies accumulating over multiple genes. Motivated by this methodological gap, we sought to develop algorithms that fuse genetic information (sets of functionally related genes) with brain data to generate functional neuroanatomical maps underlying a given brain function or behavior *in silico*.

We hypothesize that functional synergies of gene sets are best reflected in their cumulative weights on higher order features of structural (connectomes) or functional (resting state) brain networks. Based on this, we developed a method that generates functional neuroanatomical maps of functionally related gene sets from literature meta-analyses or genetic databases. We demonstrate that cumulative gene expression reflects those functional synergies. Calculating the effects of cumulative gene expression on different network measures (6, 7) proved to be sufficient for predicting functional neuroanatomy of multigenic brain functions and behavior. When applied to gene sets from genome wide association studies, quantitative trait loci (QTL) analyses or neurogenetic databases, these calculations allowed to predict brain circuits underlying complex behavioral traits in mice and human.

## Results

The method was developed on the Allen Mouse Brain Atlas (AMBA) gene expression and connectivity data framework (8, 9), a widely used mouse brain database. The mouse brain is currently the most advanced template for integrated network studies of mammalian brains with extensive gene expression and connectomic information available (8, 9). However, the method as such is general and can be applied straight forward to data from any other species such as human. The code has been optimized for low cost parallel computing.

Specifically, our method employs genetic-functional associations as inputs for weighting brain data. We fused a set of genes associated with a given brain function or behavior with gene expression maps and connectome (as structural brain network) (Fig. 1). We define the input set **T** of genes out of a genome-wide set **G**. The spatial brain gene expression data is imported pre-aligned to a common reference space from AMBA. The gene expression data consists of ordered lists of gene expression densities (10) retrieved from the AMBA for a set of spatial grid positions **D** = {d_i_}_i=1..n_ and stored as gene expression density volumes **D**(**T**) and **D**(**G**). Gene expression density is not location invariant. For example, cortical and thalamic areas have a higher mean gene expression density than the rest of the brain. Spatial bias introduced by this variance was compensated by the standardization (Z-Score) of **D**(**T**) genome-wide, such that expression density distributions at every spatial position are standard-normal distributed over **G.** Subsequently, these data sets were standardized in their spatial distribution pattern to adjust for differences between genes within the overall brain expression density.

**Figure 1.**
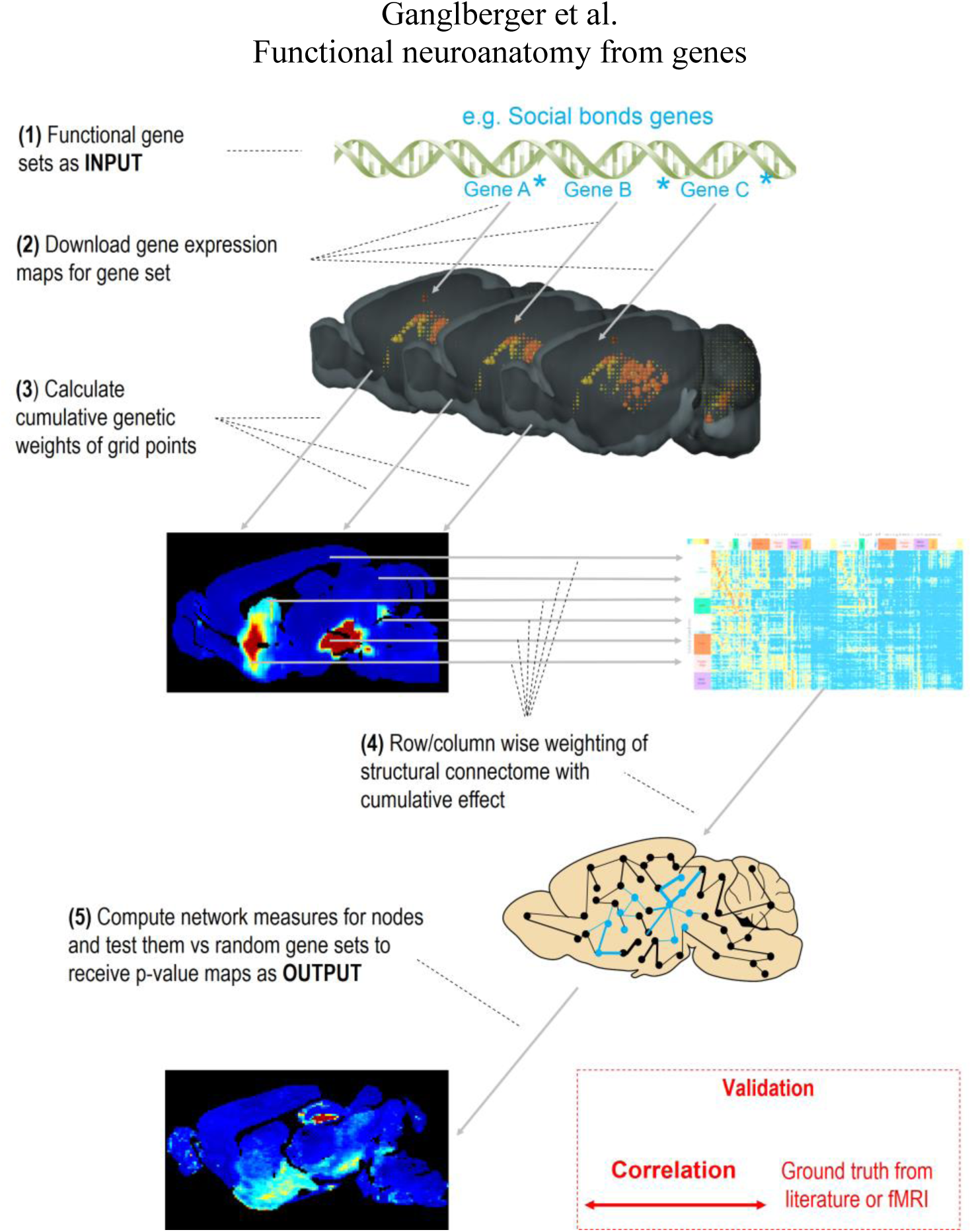
Computational workflow. A functionally-related gene set serves as input (1). For this gene set, gene expression data is retrieved (2), normalized and used to calculate a cumulative genetic effect (3). The cumulative effect is used to weight a structural connectivity matrix column or row wise (4). On the weighted network, network measures are computed and statistically evaluated by Z-tests against a null distribution (network measures based on random gene sets) (5). The output is a voxel-wise p-value map for every network measure. The results can be evaluated by computing correlation with ground truth from literature or fMRI.

Next, we sought to determine the cumulative genetic weight of **T** in **D** and calculated the synergy **S**, defined as the trimmed mean of the normalized **D** for all genes in set **T**. Trimming reduced sampling artifacts in gene density maps, like image artifacts that appear as outliers with high density scores (e.g. air bubbles) (11). The functional relation between genes and neuroanatomy is expressed by weighting either incoming or outgoing connections of every spatial sample point by **S**. Given the directed AMBA connectome as a connectivity matrix **C** ∈ **R**^n × n^ (where rows represent source regions, and columns target regions), an incoming- or outgoing-weighted connectome is defined as the row- or column-wise multiplication of **C** by **S.** To account for higher order synergies within functional maps, we computed those maps from incoming and outgoing node strengths as local network measures (12) in the weighted connectomes. For statistical evaluation, we compared the position-wise node strength measures to randomly drawn gene sets (n=1000) from the genome-wide set **G** by Z-tests (Fig. 1). We adjusted the False Discovery Rate (FDR) of the p-values with the Benjamini-Hochberg (13) method. The results in this paper are all significant under a FDR <5% (unless indicated otherwise). Ultimately, these operations generated a p-value map (a p-value for every sampling position) for every effect and brain function. To add structural context, these maps were combined (minimum p-value of effects) and projected onto the connectome, building structural networks of functionally weighted nodes that are functionally related to the input gene set. A detailed description can be found in the Supplementary Experimental Procedures.

To assess if this computational approach allows to identify function-specific brain circuitry, we focused on several well-studied gene sets, for which functional associations and functional neuroanatomy are comprehensively documented: genes associated with dopaminergic signaling, social behavior, feeding, hypothalamic–pituitary–adrenal (HPA) stress axis and synaptic plasticity. With these gene sets, we recaptured known functional neuroanatomy from literature.

For instance, genes associated with social behavior recapitulated their known functional neuroanatomy (Fig. 2*A*, Supplementary Data 1) (14–20). Similarly, we were able to pick up the functional neuroanatomy (Supplementary Data 3 Case 1-10*A,B,C*, Supplementary Data 1) for other functionally-associated gene sets (Supplementary Data 3 Case 1-10*D*) including dopamine (DA) signaling, which revealed the classical DA reward VTA-ACB pathway and also motor-related connections like SN-GP (21–24). The method allowed detecting the known feeding-related neuroanatomy based on genes associated with feeding, like orexin, neuropeptide Y (NPY), Agouti related protein (AgRP), proopiomelanocortin (POMC), melanocortin or leptin receptors (25–28). Different stress and fear/anxiety-related genes accumulate in the HPA axis, areas involved in control and regulation of stress and brain regions involved in processing fear/anxiety (29–34). We also investigated gene sets for synaptic plasticity, learning and memory. As expected, these genes highlight major sites of behavioral and functional plasticity in the brain (e.g., cortex, hippocampus, amygdala) (35–44).

**Figure 2.**
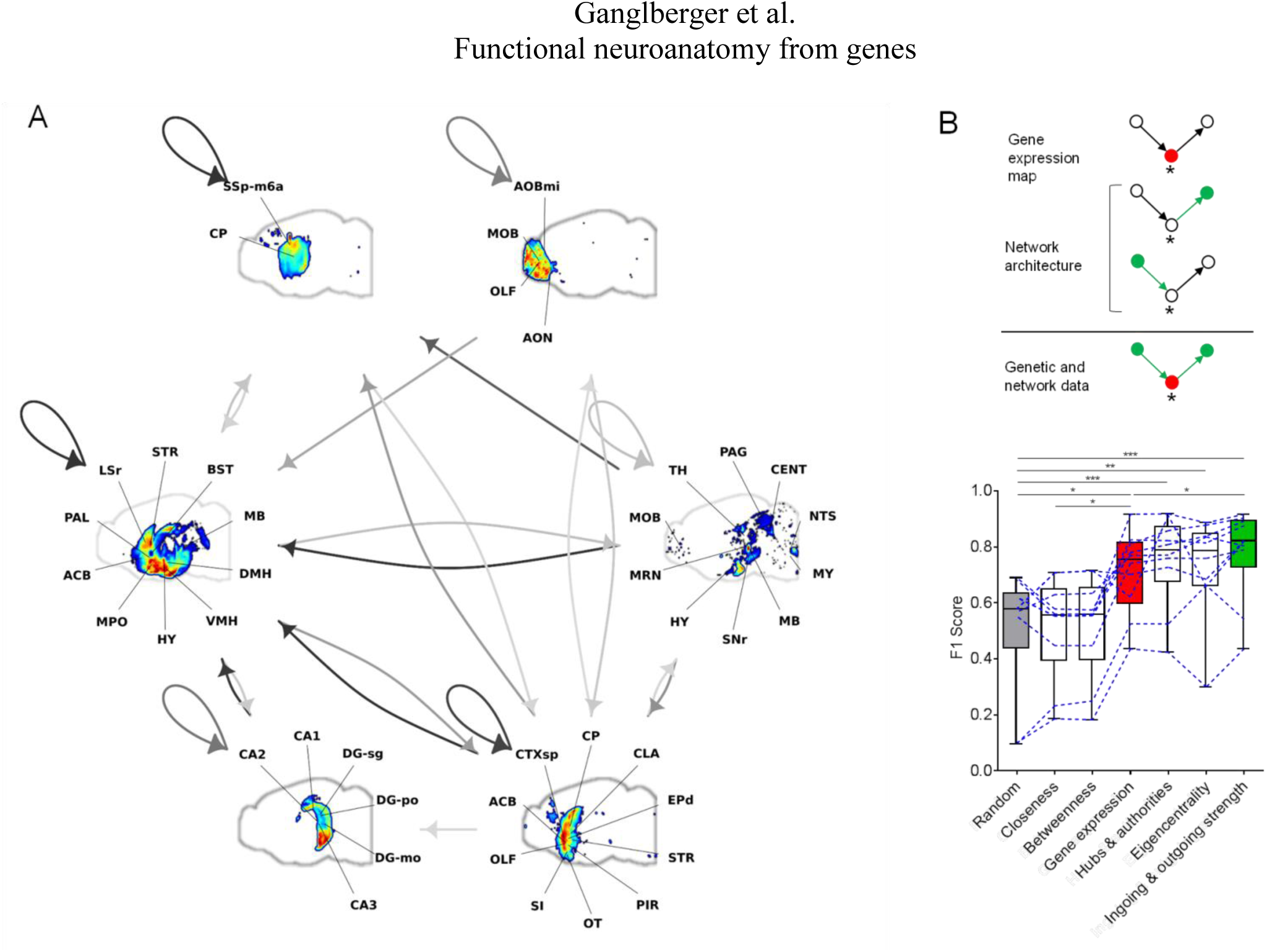
Recovery of known functional anatomy from test gene sets. (*A*) Clustered nodes of a functional anatomical map associated with a gene set for social behavior, overlayed with structural connectivity (grey arrows). The top-ranked networks include olfactory bulb (MOB), olfactory tubercle (OT), endopiriform nucleus (EPd), substantia innominata (SI), hypothalamus (HY) and hypothalamic nuclei (dorsomedial nucleus of the hypothalamus (DMH), lateral hypothalamic area (LHA, not indicated by label), ventromedial hypothalamic nucleus (VMH)), hippocampus (particularly CA2 region), midbrain (MB), including periaqueductal gray (PAG) and ventral tegmental area (VTA, not indicated by label), and nucleus accumbens (ACB). The pseudo-color scale of the nodes (colored voxels) indicates the voxel-wise accumulation of genetic weights, the intensity of the edges (arrows) the structural strength of the connection between the nodes. Loops indicate within node connections. For a complete list of abbreviations see Supplementary Tab. 1. (*B*) *Top*, Integration of first and second order network measures. The asterisk indicates a node with accumulated genetic weight. Red and green indicate sites with increased weight in first and second order measures, respectively. *Bottom*, Node-wise comparison of predicted maps to ground truth for 10 test sets. F_1_-scores increase from random classification to expression sites and to second order network measures significantly (Benjamini & Hochberg corrected One-way ANOVA on ranks, Ingoing & outgoing network strength vs Expression sites; p<0.05, Ingoing & outgoing network strength vs Random; p<0.001, Expression sites vs Random; p<0.05, Eigencentrality vs Random; p<0.01, Hubs & authorities vs Random; p<0.05). The individual F_1_ scores for each prediction are shown as dotted lines. Bars indicate median and interquartile range. Incoming & Outgoing node strength, Hubs, Authorities, Closeness, Betweenness and Eigencentrality were tested, node strength showed the highest F_1_ score.

To assess these predictions quantitatively, we collected the ground truth in form of network nodes representing regions functionally associated with these 10 gene sets from literature (Supplementary Data 2). We calculated the F_1_-score (45) of precision and recall for a binary classification of the ordered voxel-wise p-values. We used this with first order network measures (expression site; genetic weight at the node itself) and second order network measures (incoming/outgoing node strength from/to nodes with accumulated genetic weight, as well as Hub score, Authority score, Closeness, Betweenness, and Eigencentrality) (Fig. 2*B*). The computational predictions correlated significantly with the known functional neuroanatomy from literature (Fig. 2*B*, *bottom*, right bar), indicating that our method assembles meaningful functional neuroanatomical maps from genetic data.

The predictive power increased from first order measures (Fig. 2*B*, *bottom*, middle bar) to second order measures (Fig. 2*B*, *bottom*, right bar). This indicates that second order network measures detected regions not identified by gene expression alone, yet are integrated within the same neuroanatomical map. Results for node strength showed that the prediction accuracy was superior to other network measures, and is therefore sufficient for further analysis. Importantly, our approach is calculated at 100 µm voxel resolution, free from *a priori* constraints from anatomical annotations and fully compatible with small rodent MRI. Thus, it is suitable to refine structure-function relationships beyond neuroanatomical scales and has the potential to identify additional nodes and subdivisions within predefined anatomical regions with possible distinct physiological functions.

To further support our findings, we overlayed computed functional maps with those obtained experimentally with fMRI. Important in the context of this paper, pain data offers the possibility to link genetics with actual fMRI (46–48) in mice. In fact, for the pain-related gene sets (Supplementary Note 2, Supplementary Table 3 and Supplementary Data 3 Case 11-30d), the *in silico* predicted functional maps in mouse brain were reproducing large portions of the functional neuroanatomy observed with Blood-Oxygen-Level-Dependent functional magnetic resonance imaging (BOLD fMRI data, warped onto the AMBA reference space by optimized ANTS (49) parametrization) *in vivo* (Fig. 3*A*, b). This further substantiates the validity of our approach. While our method seemed to fit best with sets of >4 genes (Supplementary Fig. 2), predictions were also informative at the single-gene level. Functional imaging data of Cacna2d3 mutants, a highly conserved pain gene, revealed altered thalamo-cortical connectivity and synesthesia after thermal stimulation in mutant mice (50). The predicted maps computed from Cacna2d3 alone (Fig. 3*A*, *top right*) recaptured pain functional neuroanatomy from fMRI (Fig. 3*A*, *bottom left*, 3*B*) and pain maps that are affected by this gene (Fig. 3*A*, *bottom right*, Fig. 3*B*). Nevertheless, the single gene operations will depend heavily on the gene itself, and so we recommend to use gene sets for the most efficient and accurate functional neuroanatomy integration.

**Figure 3.**
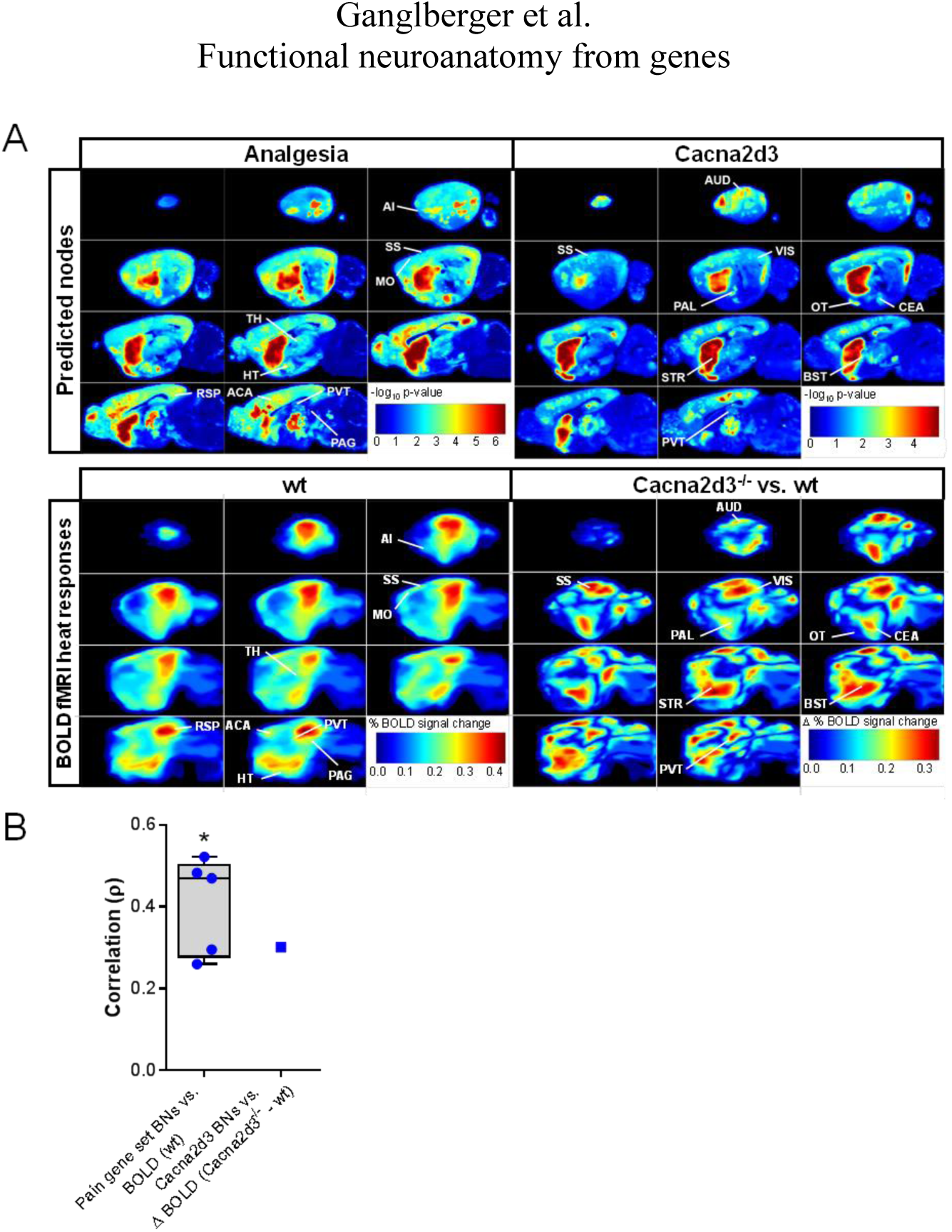
Computed functional maps correlate with BOLD fMRI of pain-related states. (*A*) Similarity of functional maps nodes predicted for analgesia gene sets and Cacna2d3 gene (*top*) to nodes with heat evoked fMRI responses (*bottom*). The highest ranked nodes include striatum (STR), paraventricular nucleus of thalamus (PVT), bed nuclei of stria terminalis (BST), pallidum (PAL), central amygdalar nucleus (CEA), sensory cortices (somatosensory areas (SS), visual areas (VIS), auditory areas (AUD)) and olfactory tubercle (OT) and correspond to those identified by fMRI. Color bars indicate −log_10_-scaled p-values (*top*) and heat stimulus responses (% BOLD signal changes) in wt animals (*bottom left*) or differences (Δ) in heat responses between Cacna2d3^−/−^ and wt animals (% BOLD signal changes in Cacna2d3^−/−^ - % BOLD signal changes in wt animals) (*bottom right*). For a detailed list of brain regions see Supplementary Tab. 1. (*B*) Voxel-wise Spearman correlations of p-value maps predicted from pain gene sets with BOLD fMRI responses. Bars indicate median and interquartile range of Spearman's ρ Wilcoxon signed rank test against ρ=0 (n=5, *W*^+^ (15) =15, *W*^−^(15) =0, *p_one-tailed_ <0.05).

Based on these results, we explored yet unknown or only partially described effector networks of behavioral traits investigated in genetic screens or association studies. One of the challenges is that behavioral traits are largely multigenic and identifying the neural circuitry through which these traits are expressed is difficult. We expanded our analysis on pain and included fear/anxiety and autism spectrum disorder (ASD) gene sets (Supplementary Note 2) from publically available databases and published meta-studies (Supplementary Table 3). In some cases, large gene sets were clustered using the DAVID platform to parcellate them into functional category-linked subsets, and so in those cases genes are not only related by the analyzed trait, but also regarding sub-functions annotated in the database. When supplied with these gene sets, our computational method extracted meaningful functional maps (Supplementary Data 3 Case 11-30). These maps, of which node-wise comparisons are in line with their functional annotation from literature, give a comprehensive representation of functional genetic synergies underlying the respective trait (Fig. 4*A*, green squares). Interestingly, we also identified nodes so far not clearly linked to investigated functions, therefore extracting potential novel functional elements (Fig. 4*A*, blue squares). These nodes might be part of the same functional network and participate in shaping the internal states of the mammalian brain.

**Figure 4.**
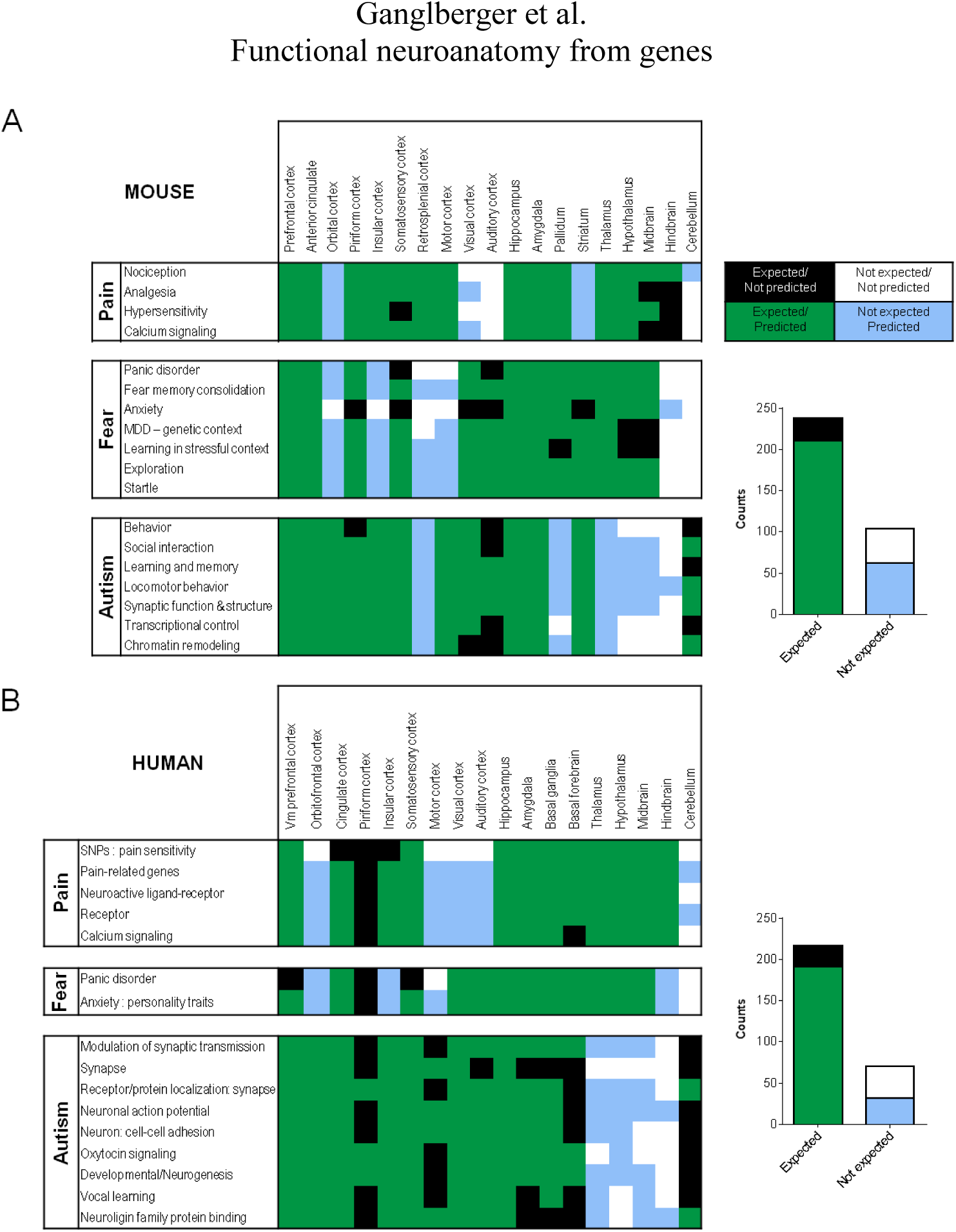
Predicting effector functional maps of behavioral traits from mouse and human genetic meta data. (*A*) *left*, Node-wise comparison of predicted mouse functional anatomy for pain, fear and autism, divided into different functional subcategories, to functional neuroanatomical annotations from literature for the top 100 p-value ranked nodes. *Right*, Quantification of the qualitative assessment. There is a significant overlap between predicted maps and functional neuroanatomical annotation (n=342; Fisher's exact test, p<0.0001). (*B*) *Left,* Node-wise comparison of predicted human functional anatomy for pain, fear and autism, divided into different functional subcategories, to functional neuroanatomical annotations from literature for the top 100 p-value ranked nodes. *Right*, Quantification of the qualitative assessment. There is a significant overlap between predicted maps and functional neuroanatomical annotation (n=288; Fisher's exact test, p<0.0001).

Extending our approach to human template based on resting state networks from fMRI (as functional brain networks) demonstrated that the methodology can be generalized to other species. Cross-validation with the meta-studies (Supplementary Data 4, Supplementary Table 2) reveals similar findings for both (Fig. 4*A*,b), demonstrating its versatility for functional exploration of the human brain in health and disease *in silico*.

## Discussion

We have developed a computational method to integrate genetic, gene expression and connectomic information from brain and genomic initiatives for rapid functional exploration of the brain *in silico*. We found that, in the brain, functionally related genes are not distributed at random but assemble into specific maps, which recapitulate functional anatomical annotations or functional data from fMRI. Cumulative effects, from expression sites alone (Fig. 2*B*, red bar), reflect functional synergies within functionally related genes, which are not directly fitted by transcriptomic similarities, usually derived from correlative analysis (Supplementary Note 1). The predictions further improved by second order network measures, which incorporate functional synergies of local gene expression that manifest in the context of higher-order interactions within the brain architecture. Incoming/Outgoing node strength (Fig. 2*B*, green bar) performed best, but not significantly better than Hubs & Authorities or Eigencentrality. This implies that nodes with the strongest effect on the network are either primary expression sites, or source/target sites thereof. Betweenness and Closeness, indicators of shortest paths in networks, outlined small distinctive nodes, that are part of functional neuroanatomy, but failed to predict the entirety of functional neuroanatomical annotations (explaining the seemingly random F_1_-score in Fig. 2*B*). The ground truth in its entirety might naturally be best explained by node strength, which reflects compounded functional synergies of regions and their afferent and efferent connections. Taken together, by fusing cumulative gene expression and best-fit network measures, we provide an optimized tool that reliably predicts functional neuroanatomical maps from genetic information.

When applied to gene sets from behavioral genetics, we demonstrated that our workflow can extract putative effector network nodes as functional brain maps which can be used to explore trait-specific circuitries. These explorations allowed to refine known functional neuroanatomy (Fig. 4, green squares). For instance, the anatomy of thalamo-cortical and cortico-cortical connections in thermal pain processing can be dissected to fine anatomical resolution (e.g., Supplementary Data 3 Case 11*E*, red arrows, note layer specificity) which could not be achieved with fMRI (Fig. 3*A*, wt). The method, based on startle response QTLs, extracted a specific and strong connection between PVT and central amygdala (Supplementary Data 3 Case 22*E*, red arrows). Interestingly this connection recently emerged as central element in fear control (51, 52). Similarly, for ASD, we identified many cortico-cortical connections (Supplementary Data 3 Case 23-29*E*, red arrows) with prediction accuracy reaching individual layers. Among similar lines, the method uncovered circuitry within regions functionally not yet associated with specific traits (Fig. 4, blue squares). For instance, the functional association of visual cortex with pain processing (53), motor cortex with startle response (54) and hypothalamic circuitry in autism (55), whose roles are understudied in the context of the respective trait or psychiatric condition, specifically at the fine anatomical or circuit level.

This can be particularly useful to pursue studies of causative role of genetic variance linked to mental diseases with unknown ethiopathology or complex course/symptomatology (with e.g., gene associations in GWAS studies as input). The method provides a holistic description of the functional neuroanatomy of a given gene set related to a meta study or behavioral trait. As such, it allows to rank order the most promising candidate regions. It has the potential to refine the functional parcellation of the brain beyond anatomical resolution, especially when performed with multiple functionally grouped gene sets at large scales. Importantly, the candidate nodes, in particular those previously not associated with those conditions, can serve as promising entry points for functional circuit dissection, e.g., with opto- and pharmacogenetic methods.

The functional relation underlying our study can be exploited to associate gene sets with specific brain functions or brain functions with specified gene sets (Supplementary Fig. 1). Importantly, our strategy applies to other neural systems (beyond mouse and human) for which genetic information, gene expression maps and connectomes are, or will be, available and allows exploration of functional brain organization in cases where actual functional data is difficult, if not impossible, to obtain.

## Acknowledgments

W. H. was supported by a grant from the European Community's Seventh Framwork Programme (FP/2007-2013) / ERC grant agreement no. 311701, the Research Institute of Molecular Pathology (IMP), Boehringer Ingelheim and the Austrian Research Promotion Agency (FFG).

### Author Contributions

F.G., J.K. and W.H. conceived the method. F.G implemented the method, performed data analysis and the quantitative validation. J.K performed the qualitative validation. J.P. and A.H. provided fMRI data. F.G., J.K., A.H., K. B. and W.H. wrote the manuscript. K.B. and W.H. jointly supervised the project.

### Competing Financial Interests

The authors declare no competing financial interests.

## Supplementary Information

**Supplementary Figures 1-2.**

**Supplementary Data 1.** P-values of first and second order effects for all cases based on region (mouse and human).

**Supplementary Data 2.** Ground truth generated from literature.

**Supplementary Data 3.** Functional neuroanatomical maps, significant regions and network visualization of all cases used in this paper for mouse.

**Supplementary Data 4.** Significant regions of all cases used in this paper for human.

**Supplementary Table 1.** Anatomical abbreviations.

**Supplemental Experimental Procedures**

### Mouse Data

The mouse connectome was retrieved as (structural) connectivity from all 2173 available injection sites (state March 2016) to their target sites given as image data, detailing projections labeled by rAAV tracers via serial two-photon tomography (9). Those sites are added up to a connectivity matrix which covers about 15 percent of the right hemisphere as source regions, and about 100% as target regions. The AMBA connectome (right hemisphere injections) was mirrored onto (left hemisphere) AMBA gene expression data. In order to also take weak connections into account, the connectome was binarized by a threshold according to Oh, S. W. *et al.* (9), Extended Data Figure 7, that minimizes the amount of false positive connections. The gene expression density is interpolated to a 100 micron resolution to match the resolution of the connectome. A Matlab script for downloading the gene expression for **T** and for **G**, as well as the AMBA connectome is provided on request.

### Human data

Gene expression by region retrieved from the Allen Human Brain Atlas (56). The Allen Institute provides an affine transformation to MNI152 (57) space by its API. We used resting state functional connectivity from the Human Connectome Project (58), which is also in MNI152space (57).

### Mathematical description

<u>Input</u> data is a functionally related gene set, more precisely a certain brain function or behavioral trait represented as a list of genes. Spatial gene expression and connectomic data were retrieved from AMBA.

<u>Data retrieval</u> was performed via the AMBA API. It allows the download of 3D spatial gene expression patterns(8) for available genes at given grid positions with a resolution of 200 microns.

We retrieve for n grid positions 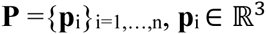 and each available gene g in the mouse genome **G** ={g_j_}_j=1..m_ (or at least a random drawn subset) the gene expression density

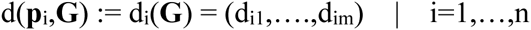

and store it as gene expression density volume

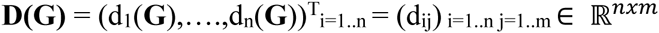

This is also done for the gene function/trait associated gene set **T** = {t_k_}_k=1,…,1_ being a subset of **G**, resulting in the expression density volume 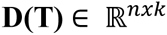.

<u>Normalization</u> of the function/trait specific expression density volume **D(T)** is performed over the genomic as well as over the spatial domain. At first, standardization in the genome space is performed so that every spatial sample point has a distribution of gene expression densities with a mean of 0 and a standard deviation of 1 over the whole genome **G**

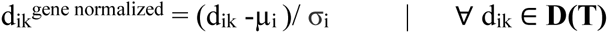

where µ_i_ = µ((d_ij_) _j=1,..,m_) and σ_i_ = σ(d_i_((d_ij_) _j=1,..,m_), d_ij_ ∈ **D(G)**.

This normalization compensated for spatial bias in the mean **density**. For example, the cerebral cortex and thalamic areas have a higher mean **density** than the rest of the brain.

In a second stage, standardization is performed for **D^gene normalized^** (T) = (d_ij_^**gene normalized**^) in the spatial domain, so that each gene in **T** has a distribution of gene expression densities with a mean of 0 and a standard deviation of 1 over all sample positions

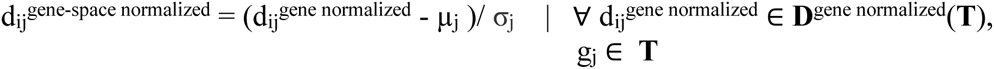

where µ_j_^gene normalized^ = µ (d_ik_^gene normalized^)_k=1,…,l_) and σ_j_^gene normalized^ = σ (d_ik_^gene normalized^)_k=1,…,l_), d_ik_ ∈ **D(T)**. We replaced missing values with 0 (which is the most likely value that a value can have after normalization in genome space) for the calculation of µ_j_ and σ_j_ to compensate for missing lateral slices from AMBA.

<u>Effect calculation</u> is based on the trimmed mean of the gene-space normalized densitiy of all genes in the function/trait set, that is called synergy **S** = (s(p_i_))_i=1..n_ where

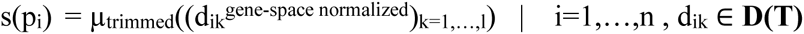

With the synergy **S**, several effects can be computed. Effects are divided into first order and second order effects:

<u>First order effects</u> do not take the context of the network into account. The synergy **S** is a first order effect itself, since **S** represents the gene function/trait association of every sample point. Other first order effects would be the µ((d_ik_^gene-space normalized^)_k=1,…,i)_ (which is not robust to image artifacts like bubbles), or max((d_ik_^gene-space normalized^)_k=1,…,l_),((d_ik_^gene-space normalized^)_k=1,…,l_)

<u>Second order effects</u> show the influence of the function/trait in the context of the network. The function/trait-network association is expressed by weighting either incoming or outgoing connections of every sample position by **S**, depending on the scope of interest (afferent or efferent connections). Given a directed connectome as connectivity matrix

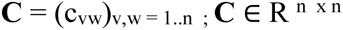

where the rows represent the source regions, the columns target regions, either an incoming **C^weighted in^** or outgoing **C^weighted out^** weighted directed connectome is defined as

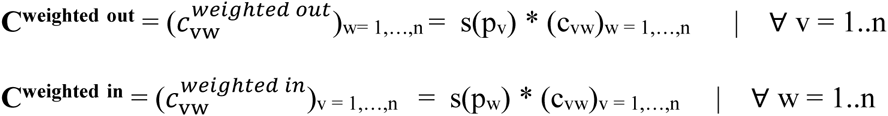

The second order effects on the network are computed by local network measures such as incoming/outgoing node strength, hubs, authorities, closeness, betweenness and eigencentrality on both incoming and outcoming weighted connectomes **C**^weighted in^ and **C**^weighted out^. We showed in Fig. *2B*, that incoming/outgoing node strength performed best on predicting our test data and is therefore stated exemplary. The incoming node strength (sum of incoming connections for every node) of **C**^weighted in^ and **C**^weighted out^ is defined as

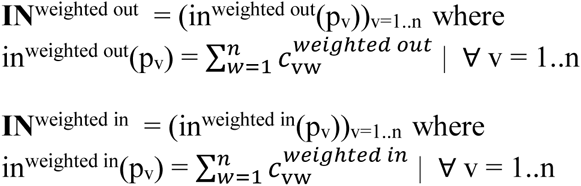

and the outgoing node strength (sum of outgoing connections for every node) as

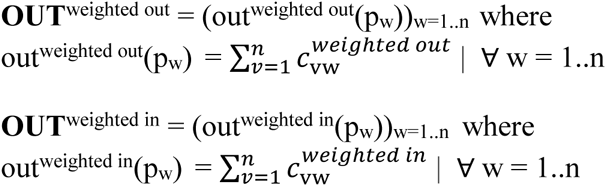

<u>Statistical evaluation</u> of the computed effects (first and second order) are performed by comparing them to the effects of random drawn gene sets (genome-wide randomized function/trait-gene association) from the genome **G**.

1. Calculate the network effects for a function/trait **T**.
2. Draw >=1000 random set of genes from the genome **G** with equal size of **T**.
3. Calculate the first and second order effects for every random set.
4. P-values for the effects of **T** can be computed for every spatial sample position by performing a Z-test against the null-distribution represented by the >=1000 random effects since every spatial sample point is normally distributed in the gene dimension (verified by KS tests).

The significance of **IN**^weighted out^ can be interpreted as nodes that are receiving from primary expression sites (regions with high **S)**, while **OUT**^weighted in^ shows regions projecting to primary expression sites. P-value calculations of **IN**^weighted in^ and **OUT**^weighted^ out are numerically equal to the p-value calculation of **S** (for a node degree>0), since for those cases the sum of incoming and outgoing connections are constant factors when compared to random effects. We point this out to clarify the p-value calculation of **IN**^weighted in^ and **OUT**^weighted out^ can be substituted by **S** for computational reasons.

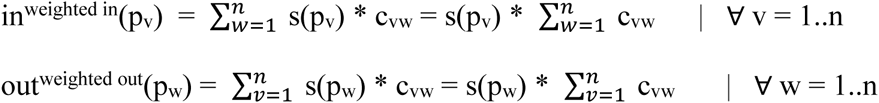

Due to the multiple comparison problem, we adjust the FDR of the p-values of the effects by the Benjamini-Hochberg (13) method. The results in this paper are all significant under a FDR <5% (if not indicated otherwise).

<u>Output</u> is a p-value map (a p-value for every spatial sample point) for every effect. In this paper, **S**, **IN, OUT** are used due to their fast computation, simplicity and biological significance.

### Code availability

The code for retrieving data (gene expression, mouse connectome) from the AMBA API consists of a Matlab script whose single input parameter is a .csv with function/trait information as a list of gene symbols and Entrez IDs. The main algorithm was implemented as an R-script that uses the generated files (downloaded data from AMBA) of the Matlab script to normalize, calculate and carry out a statistical evaluation to generate p-value maps and structural network visualization for every testcase. The statistical evaluation, which was randomized because of the extent of the computational task, is parallelized.

MATLAB- and R-codes will be made publically available under an open source license for non-commercial use upon acceptance of the paper for publication.

### Figure generation

Figures were generated with a R-script that will be provided on request. It uses the p-value maps of the method to generate slice-views of different effects, heatmaps with statistical measures of the effects and gene expression, clustered networks, csv-files with raw data and precision-recall heatmaps (for data with ground truth).

<u>Slice-views</u>: Slice-views show 11 maximum intensity projections of 5 sagittal slices each of a 132x80x114 voxel volume (which represents spatial sample positions) that shows the left hemisphere of the mouse brain. Slice-views are used to visualize a log-scaled mapping of first order p-values (of **S**), second order incoming node strength **IN** (regions that are targets of first order regions) and second order **OUT** (regions projecting to first order regions). At the bottom-right corner is a color-bar, indicating the minus log_10_scaled p-values, the threshold for false positive FDR (10% solid line, 5% dotted line). Slice-views of all testcases can be found in Supplementary Data 3 Case 1-30*A, B, C*.

<u>Heatmaps</u>: Heatmaps in Supplementary Data 3 Case 1-30*D* and Supplementary Data 4 show the log-scaled p-values of first and second order effects as well as single gene effects (gene expression density of a gene vs gene expression density of the genome) for every significant region (a region that has at least one voxel with significant first or second order effect). The regions are color-coded (on the left side) corresponding to the AMBA, and given by their acronym on the right side. Similar information can be found in the attached csv files (Supplementary Data 1) which contain the region-wise p-values of first and second order effects.

<u>Clustered network graphs</u>: We clustered our test sets via hierarchical clustering with Ward's Criterion (59) using the R function `hclust(*, "ward.D2")`. To ensure that voxels with similar connections are within the same cluster, they are clustered by their Pearson-correlation coefficient of their connectivity. To visualize the clusters, we plotted a sagittally-projected heatmap of their combined p-value (minimum p-value of effects), surrounded by labels. The connectivity between clusters is shown by the sum of connectivity (normalized by injection volume) between the clustered regions given as grey-scale. All graphs can be found in Fig. 2*A* and Supplementary Data 3 Case 1-30*E*.

<u>F_1_-score bar-chart</u>: Based on available ground truth from the literature (Supplementary Data 2), we calculated the F_1_-score (45) based on the precision and recall for a binary classification of ordered p-values. It doesn't take the true negative rate into account, which is acceptable for the following reason: The literature-based ground truth is region based. This means we can identify

- true positives (a positive classified voxel within a region of the ground truth)
- false positive (a positive classified voxel outside a region of the ground truth)

but not

- true negative (a negative classified voxel outside a region of the ground truth), since the total set of regions of the functional neuroanatomy are still unknown
- false negatives (a negative classified voxel within the ground truth), since it is possible that only a subset of the ground truth region is specific for functional neuroanatomy.

For the calculation of the F_1_-score, respectively precision and recall, the precision is computed as the ratio of true positive voxels to the amount of positive voxels. For a voxel-based recall, a false negative rate would be necessary, and so we used the region-based recall, the ratio of positive classified regions to ground truth regions. We defined a positive classfied region if at least 5% of the voxels of a region is positive (to account for noise). P-value maps for the F_1_-score bar chart were computed at 200 micron resolution due to extensive computational network measures.

### Technical resources

We used the Amazon elastic cloud computing service with an "r3.8xlarge" instance (32 cores, 244 GB RAM) (60). More than 100 GB RAM is recommended, 40 GB alone to hold the connectivity matrix in the memory. Additional memory is needed for parallel processing (approximately 3 GB per core). We tested the R-scripts with 30 cores. The computation uses about 200 GB Ram and takes between 1 and 2 hours per testcase (depending on the amount of genes in a set) to calculate the p-values for first and second order effects. The clustering for the circle-graphs are also parallelized. Depending on the size of the significant areas, clustering takes between 30 minutes to 3 hours.

### General statistics

Unless indicated otherwise, data were tested for normality by Kolmogorov–Smirnov or D'Agostino & Pearson tests at α<0.05 and analyzed non-parametrically if tests didn’t pass. Predicted functional neuroanatomy maps were compared to ground truth from fMRI using a Spearman correlation of the −log_10_-scaled voxel-wise p-value of predicted nodes, set to p=10^−3^ for all p<10^−3^, to BOLD heat responses of wt animals or differences in BOLD heat responses in Cacna2d3 mutant vs. wt animals, respectively. To compensate for registration errors between the AMBA reference space and fMRI data, these comparisons were performed on volumes downsampled to 400 μm spatial resolution.

## Supplementary Note 1

Investigating functional and structural brain network data and its analysis is an ongoing challenge (61). Bullmore and Sporns (61) described the exploration of structural and functional brain networks as a multi-stage approach, beginning with the separate creation of structural and functional connectivity matrices based on anatomical parcellations. Network measures, such as node degree, node strength, hubs, centrality, betweenness etc., indicate network properties of interest when compared to equivalent measures of a population of random networks (null-distribution). A local (region-wise) or global (Mantel-test) (62) comparison reveals functional and structural correspondences of the networks.

The integration of genetic information facilitates insight into the influence on neuronal activity and structural organization of the brain (1). French and Pavlidis (1) compared cortical and subcortical regions of a rat connectome (63) and AMBA gene expression data (8) using Spearman's rank correlation to show that brain regions with similar expression patterns have more similar connectivity profiles. The similarities are close enough that a computational model by Ji et al (64) could predict structural connectivity by gene expression profiles. 4048 genes with coronal spatial expression data were used as individual features in a sparse model to obtain a predictive accuracy of 93% on anatomical parcellations. A follow up study proved that this also works on mesoscale-resolution (voxels at 200 micron resolution) (65).

A combined approach of comparing structural connectivity, gene co-expression correlation and functional networks was investigated by (3). Resting-state fMRI networks (default-mode, salience, sensorimotor and visuospatial) were used as a starting point to identify functionally related cortical regions in mice and humans. The strength fraction (scaled node strength of gene co-expression networks) between those regions was significantly more similar than to the remaining brain regions (tested by permutation tests). Genes that are related to the four functional networks were identified by ranking them by their marginal influence on the strength fraction. A gene co-expression matrix including only top-ranked genes was compared to structural connectivity using the Mantel procedure (62) and were significant compared to a sample of 10,000 random gene sets. (2) used Spearman's rank correlation between node degree of structural connectivity and gene co-expression of gene sets related to Gene Ontology groups (cellular composition and biological process) to assess how structural connectivity is genetically driven. Connectivity related Gene Ontology groups were also used by Fulcher and Fornito (66). They showed that the mean gene co-expression correlation of groups related to biological processes are higher for connections involving structural “hubs” (node degree over threshold) vs non-hubs indicates topological specializations of interregional connections. Structural network hubs were also found to correspond to known functional networks from the literature (4, 5). Compared to other studies (1–3, 66) which used node strength or variations of it, Rubinov and Sporns (12) assessed other structural network parameters, such as community structures, hierarchical modules, high-low cost sub-networks etc.

An overview of related work and its modalities can be found in Supplementary Table 1. Apart from Fakhry and Ji (65), who used high-resolution prediction, the studies cited were computed on anatomically parcellated mouse brains (Richiardi and Altmann (3) also used human data). Our approach was performed on 100-micron grid parcellation. In contrast to Richiardi and Altmann (3), where functionally related gene sets were products of their marginal influence on resting-state networks, we used functionally-linked gene sets as the entry point of our method. Fulcher and Fornito, as well as French et al. (2, 66) showed the influence of Gene Ontology groups of biological processes on structural networks, while our approach utilized sets from gene association studies (database-mining, QTL analyses or SNPs) and that can be directly linked to certain behavioral or mental features. Known functional networks from the literature confirmed our results as well as the correlation with resting state fMRI.

Comparing gene co-expression correlation to structural connectivity is a common approach for assessing brain structures with genetic functionality (1–3, 64–66, 4, 5). The novelty in our paradigm is weighting structural connectivity with functionally related, cumulative gene. It is not only comparing networks, but it shows the direct effect of functionally related gene expression on brain anatomy. Those effects were encountered by node strength, which we proved to be a sufficient indicator, but also with various other network measures.

## Supplementary Note 2

Pain sensation is biomedically one of the most important brain functions. While physiological sensation is essential to protect the organism and to avoid harm, it is very often a result of diseases or pathological/abnormal processes when the sensory information does not reflect the factual danger from the environment. Pain gene sets from mice and human were taken from literature and databases (Supplementary Data 2) (67, 68). pre-clustered or pre-assigned to subcategories based on behavioral phenotype (nociception, analgesia, hypersensitivity) or functional annotations (Gene Ontology (GO)), calcium signaling = calmodulin binding+calcium ion transport associated genes related to pain processing). For the human case we chose a metastudy combining SNPs associated with pain sensitivity or we extracted subcategories (obtained using the DAVID platform based on functional annotation) from the database for pain-related genes. We also used the Calcium signaling category as a set based on evolutionary conserved pain genes. Importantly, the effector networks from most of these gene sets could be linked to known pain-related areas in the brain (46, 48, 69, 70), but also other regions such as piriform and entorhinal cortices, nucleus accumbens and VTA (Fig. 4*A*). Functional neuroanatomy maps from these gene sets, and the single gene Cacna2d3, were also compared to fMRI pain responses of wt and mutant animals, respectively (50) (Fig. 3*A*). The maps derived from the gene sets (except nociception) were similar to the expected pain network from the mouse fMRI (Fig. 3*A*). The Cacna2d3-dependent maps identified by our method retraced Cacna2d3's functional genetic effects on pain processing in fMRI in regions like striatum, olfactory areas, somatosensory cortex, hippocampus, hypothalamus, paraventricular nucleus of thalamus (PVT) and basal ganglia. Similarly, for the human gene sets (Fig. 4*B*), we obtained the brain regions known to be involved in pain processing, including central grey, PVT, insular and somatosensory cortex, but also VTA – as in the mouse case – or higher order associative cortices which are responsible for self-awareness and conscious perception of pain.

Fear and anxiety-related genes were retrieved from JAX QTLs database (mouse) or from literature (mouse and human) (71, 72), pre-assigned to behavioral phenotypes (startle response, exploration, anxiety, depression and panic disorder). Again, the computed maps (mouse and human) contained nodes with a fitting functional annotation, like fear-related regions in the amygdalar complex, prefrontal cortex, thalamic or midbrain structures (73–78). Moreover, the main nodes detected by our method are in line with their associated functional subcategory, e.g. startle behavior was linked to insular cortex and PVT, while mental disorders were linked to insular cortex, ACB and VTA (Fig. 4*A*). For the panic disorder category, we can see differences in cortical regions identified for mouse and human. For example, human data, unlike the mouse, lacks vmPFC, somatosensory or motor corices, while we did not detect the auditory cortex in the mouse brain (Fig. 4).

For autism-related genes, we retrieved 183 genes implicated in behavioral phenotypes in mouse models of ASD and 739 autism-associated genes in humans from Autdb database (79) and clustered the genes with DAVID (80), for further analysis, we chose functional annotation categories that were the most relevant for ASD modeling: linked to behavior, cognitive abilities, synaptic functions and cellular level processes. Similar to the other gene sets, the computationally predicted maps contained nodes related to autistic brain function (71, 81–88), in the case of the human brain several cortical, subcortical and cerebellar areas were not identified (Fig. 4*B)*.

To sum up, we were able to identify most of the known functionally involved brain regions for all of the investigated categories based on mouse and human data. Additionally, for different specific subcategories the method identified functionally relevant structures which were found at the highest positions in rank-order lists. Taking together all the data, the method can also be a useful tool for identifying novel functional targets, potentially involved in traits linked to the genetic input. With this, we can bridge already known functional systems using potential new-still unexplored - connections or even identify new functional networks. For more detailed information please see Supplementary Data 1, 2, Fig. 3, Supplementary Data 3 Case 11-29 (for mouse) and Supplementary Data 4 (for human).

**Supplementary Table 2.**
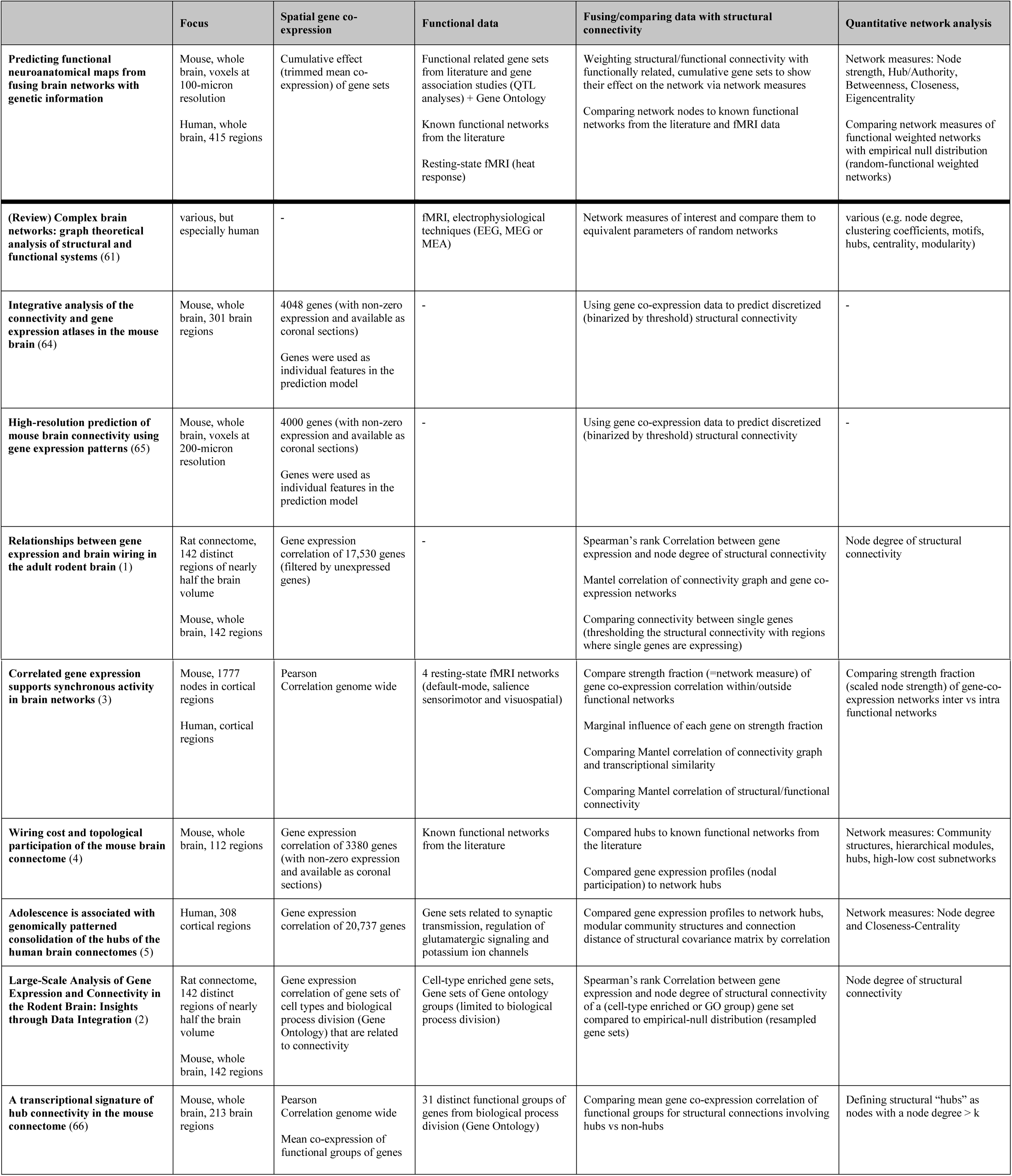
Outline of related work with focus on the quantitative analysis of networks that are either functional, structural, derived from gene expression, or a combination thereof.

|  | Focus | Spatial gene co-expression | Functional data | Fusing/comparing data with structural connectivity | Quantitative network analysis |
| --- | --- | --- | --- | --- | --- |
| **Predicting functional neuroanatomical maps from fusing brain networks with genetic information** | Mouse, whole brain, voxels at 100-micron resolution<br>Human, whole brain, 415 regions | Cumulative effect (trimmed mean co-expression) of gene sets | Functional related gene sets from literature and gene association studies (QTL analyses) + Gene Ontology<br>Known functional networks from the literature<br>Resting-state fMRI (heat response) | Weighting structural/functional connectivity with functionally related, cumulative gene sets to show their effect on the network via network measures<br>Comparing network nodes to known functional networks from the literature and fMRI data | Network measures: Node strength, Hub/Authority, Betweenness, Closeness, Eigencentrality<br>Comparing network measures of functional weighted networks with empirical null distribution (random-functional weighted networks) |
| **(Review) Complex brain networks: graph theoretical analysis of structural and functional systems (61)** | various, but especially human | - | fMRI, electrophysiological techniques (EEG, MEG or MEA) | Network measures of interest and compare them to equivalent parameters of random networks | various (e.g. node degree, clustering coefficients, motifs, hubs, centrality, modularity) |
| **Integrative analysis of the connectivity and gene expression atlases in the mouse brain (64)** | Mouse, whole brain, 301 brain regions | 4048 genes (with non-zero expression and available as coronal sections)<br>Genes were used as individual features in the prediction model | - | Using gene co-expression data to predict discretized (binarized by threshold) structural connectivity | - |
| **High-resolution prediction of mouse brain connectivity using gene expression patterns (65)** | Mouse, whole brain, voxels at 200-micron resolution | 4000 genes (with non-zero expression and available as coronal sections)<br>Genes were used as individual features in the prediction model | - | Using gene co-expression data to predict discretized (binarized by threshold) structural connectivity | - |
| **Relationships between gene expression and brain wiring in the adult rodent brain (1)** | Rat connectome, 142 distinct regions of nearly half the brain volume<br>Mouse, whole brain, 142 regions | Gene expression correlation of 17,530 genes (filtered by unexpressed genes) | - | Spearman's rank Correlation between gene expression and node degree of structural connectivity<br>Mantel correlation of connectivity graph and gene co-expression networks<br>Comparing connectivity between single genes (thresholding the structural connectivity with regions where single genes are expressing) | Node degree of structural connectivity |
| **Correlated gene expression supports synchronous activity in brain networks (3)** | Mouse, 1777 nodes in cortical regions<br>Human, cortical regions | Pearson<br>Correlation genome wide | 4 resting-state fMRI networks (default-mode, salience sensorimotor and visuospatial) | Compare strength fraction (=network measure) of gene co-expression correlation within/outside functional networks<br>Marginal influence of each gene on strength fraction<br>Comparing Mantel correlation of connectivity graph and transcriptional similarity<br>Comparing Mantel correlation of structural/functional connectivity | Comparing strength fraction (scaled node strength) of gene-co-expression networks inter vs intra functional networks |
| **Wiring cost and topological participation of the mouse brain connectome (4)** | Mouse, whole brain, 112 regions | Gene expression correlation of 3380 genes (with non-zero expression and available as coronal sections) | Known functional networks from the literature | Compared hubs to known functional networks from the literature<br>Compared gene expression profiles (nodal participation) to network hubs | Network measures: Community structures, hierarchical modules, hubs, high-low cost subnetworks |
| **Adolescence is associated with genomically patterned consolidation of the hubs of the human brain connectomes (5)** | Human, 308 cortical regions | Gene expression correlation of 20,737 genes | Gene sets related to synaptic transmission, regulation of glutamatergic signaling and potassium ion channels | Compared gene expression profiles to network hubs, modular community structures and connection distance of structural covariance matrix by correlation | Network measures: Node degree and Closeness-Centrality |
| **Large-Scale Analysis of Gene Expression and Connectivity in the Rodent Brain: Insights through Data Integration (2)** | Rat connectome, 142 distinct regions of nearly half the brain volume<br>Mouse, whole brain, 142 regions | Gene expression correlation of gene sets of cell types and biological process division (Gene Ontology) that are related to connectivity | Cell-type enriched gene sets, Gene sets of Gene ontology groups (limited to biological process division) | Spearman's rank Correlation between gene expression and node degree of structural connectivity of a (cell-type enriched or GO group) gene set compared to empirical-null distribution (resampled gene sets) | Node degree of structural connectivity |
| **A transcriptional signature of hub connectivity in the mouse connectome (66)** | Mouse, whole brain, 213 brain regions | Pearson<br>Correlation genome wide<br>Mean co-expression of functional groups of genes | 31 distinct functional groups of genes from biological process division (Gene Ontology) | Comparing mean gene co-expression correlation of functional groups for structural connections involving hubs vs non-hubs | Defining structural "hubs" as nodes with a node degree > k |

**Supplementary Table 3.** Summary of mouse and human functional genetic data collection.

|  |  | MOUSE | HUMAN |
| --- | --- | --- | --- |
| Pain | Meta-study:<br>Database | Functional classes according to the Pain database (67) | Common human genetic variants (89–91) + Functional analysis with DAVID (80) |
|  | Meta-study:<br>literature screen |  | SNPs (89) |
|  | Experimental<br>data (literature) | Neuronal-specific RNAi knock-down strategy in adult *Drosophila* (68) + Functional clustering |  |
| Fear | Meta-study:<br>Database | Functional classes from JAX database: QTLs (72) |  |
|  | Meta-study:<br>literature screen | Multidisciplinary integration of human (71, 90) and mouse (71, 91, 92, 93, 94) data |  |
| Autism | Meta-study:<br>Database | Collection of all genes connected to ASD in humans and relevant animal models (AutDB) (79) + Functional analysis (DAVID) (80) |  |

